# Structures of PKA-phospholamban complexes reveal a mechanism of familial dilated cardiomyopathy

**DOI:** 10.1101/2021.11.16.468845

**Authors:** Juan Qin, Jingfeng Zhang, Lianyun Lin, Omid Haji-Ghassemi, Zhi Lin, Kenneth J. Woycechowsky, Filip Van Petegem, Yan Zhang, Zhiguang Yuchi

**Author notes:** Correspondence and requests for materials should be addressed to Z.Y.

## Abstract

Several mutations identified in phospholamban (PLN) have been linked to familial dilated cardiomyopathy (DCM) and heart failure, yet the underlying molecular mechanism remains controversial. PLN interacts with sarco/endoplasmic reticulum Ca^2+^-ATPase (SERCA) and regulates calcium uptake, which is modulated by the protein kinase A (PKA)-dependent phosphorylation of PLN during the fight-or-flight response. Here, we present the crystal structures of the catalytic domain of PKA in complex with wild-type and DCM-mutant PLNs. Our structures, combined with the results from other biophysical and biochemical assays, reveal a common disease mechanism: the mutations in PLN reduce its phosphorylation level by changing its conformation and weakening its interactions with PKA. In addition, we demonstrate that another more ubiquitous SERCA-regulatory peptide, called another-regulin (ALN), shares a similar mechanism mediated by PKA in regulating SERCA activity.

**Significance:** Dilated cardiomyopathy (DCM) is a common type of heart disease. Familial DCM is associated with mutations on phospholamban (PLN), but the mechanism remains elusive. Phosphorylation of PLN is known to influence its physiological function. We hypothesize that the connection between such mutations and DCM may involve decreased PLN phosphorylation levels due to less efficient binding to protein kinase A. We utilize x-ray crystallography, SPR, enzyme kinetic assays, thermal melt assays, and NMR to examine the structural and energetic consequences for PKA-catalyzed phosphorylation of PLN variants containing DCM-associated mutations. Our results provide a foundation to understand the general working mechanism of PKA and the physiological regulation of PLN by PKA, and also provide important insight into the pathological mechanism of DCM.

## Introduction

Dilated cardiomyopathy (DCM) is the most common type of cardiomyopathy, characterized by an enlarged heart with a decreased ejection fraction. It is a major cause of heart failure (*1*), affecting 40 million people globally (*1–3*). 25-35% of DCM cases have familial origin (*4*), caused by inherited mutations in genes encoding proteins involved in muscle contraction and calcium handling, including phospholamban (PLN) (*5–8*).

To initiate cardiac muscle contraction, an action potential depolarizes the sarcolemma and activates the voltage-gated calcium channel, Ca_V_1.2, which mediates Ca^2+^ influx. The small increase in the cytosolic Ca^2+^ concentration causes larger-scale calcium-induced calcium release (CICR) from the intracellular sarcoplasmic reticulum (SR) stores through cardiac ryanodine receptors (RyR2) (*9, 10*). The resulting increase of the cytosolic [Ca^2+^] from 100 nM to 10 μM induces muscle contraction. To relax the cardiac muscle, sarcoplasmic reticulum Ca^2+^-ATPases (SERCAs) on the SR membrane couple ATP hydrolysis to the pumping of Ca^2+^ back into the SR (*11, 12*). The rate and duration of SERCA-mediated SR calcium restoration affect the SR calcium load, which regulates the rate of muscle relaxation and the intensity of the next contraction.

PLN, an important regulator of SERCA, is a 6.2 kDa single-pass integral membrane protein that can reversibly inhibit SERCA activity by physically interacting with the calcium pump and thus regulate the contraction of cardiac muscle (*13–17*). The transmembrane domain of PLN interacts with SERCA via a conserved sequence motif (*18*). Inhibition of SERCA by PLN responds to changes in the free calcium concentration and redox environment of the cytosol. The potency of SERCA inhibition also depends on certain structural properties of PLN, such as its oligomerization state and phosphorylation levels. PLN can be phosphorylated by cAMP-dependent protein kinase A (PKA) and calmodulin-dependent protein kinase II (CaMKII) (*13, 16, 19*). During the fight-or-flight response, the activation of β-adrenergic receptor leads to the activation of PKA, which in turn phosphorylates a number of downstream targets regulating cardiac muscle contraction, including Ca_V_1.2, RyR2, troponin, and PLN (*20*). PKA mediates phosphorylation of PLN at Ser16, which relieves its inhibition of SERCA, increasing muscle contractility and relaxation rate (*13, 16, 21, 22*).

To date DCM mutations associated with PLN include R9C, R9H, R9L, ΔR14, R14I and I18T (*23–28*). These mutations cluster in a small “hotspot” region that has little direct contribution to the inhibition of SERCA activity (*29*). However, its neighborhood contains the critical PKA (*13, 16, 22*) and CaMKII (*21, 30*) phosphorylation sites, Ser16 and Thr17 respectively, suggesting a connection between altered regulation of phosphorylation and DCM phenotype. Among them, the R9C and ΔR14 mutations have the highest frequency and are associated with the most severe DCM phenotype (*23, 25*). R9C PLN has been studied extensively, but its DCM-causing molecular mechanism remains controversial. The initial study by Schmitt et al. suggests that compared to wild type (WT) PLN, the R9C mutant interacts more tightly with PKA, prevents dissociation of PKA from R9C PLN, and thus locks it in an inactive state, which prevents the phosphorylation of PLN at Ser16 (*23*). Ha et al. show that the R9C mutation stabilizes the pentameric form of PLN by introducing inter-subunit disulfide bonds under oxidizing conditions, decreasing inhibition of SERCA (*19*). Other hypotheses highlight the importance of altered PLN conformation (*31*), hydrophobicity (*32*) and PLN-membrane interactions (*33*) caused by the mutation. The impacts of other DCM mutations have been confirmed by genetic (*26*) or animal studies (*25*), but their disease mechanisms are far from clear.

ALN is a newly identified protein which possesses a transmembrane domain that shares the conserved SERCA-interacting sequence motif with PLN (*34*). Unlike PLN, which is specifically expressed in cardiac muscle, ALN is ubiquitously expressed in many tissues including atria and ventricle but with the highest expression levels in the ovary and testis (*34*). Although PLN and ALN diverge significantly in their cytoplasmic domains, a putative PKA recognition motif is present in ALN, suggesting the possibility that ALN is a substrate of PKA. Consistent with this notion, phosphorylation of Ser19 was detected by mass spectrometry of mouse ALN extracted from various organs (*35, 36*). However, it remains to be investigated whether this phosphorylation is carried out by PKA and whether this phosphorylation regulates the interaction of ALN with SERCA.

Here we report three crystal structures of the PKA catalytic domain (PKAc) in complex with three peptide variants of PLN, WT, R9C, and A11E. The long sought after structure of the PKAc-R9C PLN complex is critical for understanding the disease mechanism of familial DCM. Compared to the PKAc-WT PLN structure, the replacement of Arg by Cys abolishes an important electrostatic interaction, resulting in a significant conformational change of PLN and significantly reduced interactions between the two proteins. The binding affinities of various PLN peptides to PKAc were measured by surface plasmon resonance (SPR) and compared to each other. Consistent with our crystal structures, upon R9C mutation, the binding affinity to PKAc was significantly reduced compared to WT PLN. The kinetic constants of PKAc-catalyzed phosphorylation of PLN peptides were also determined and a significantly lower *k_cat_*/K_M_ was observed for R9C PLN. Our data also support the idea that other PLN mutations in the neighborhood, including DCM-related mutations at the 9th, 14th and 18th positions, share a common disease mechanism related to reduced PKA phosphorylation. The solution-phase structures of free PLN variants determined by NMR show that, individually, phosphorylation of the WT peptide and DCM-related mutations cause PLN to become more rigid, which might also contribute to the reduction of phosphorylation. We also confirm that ALN can be phosphorylated by PKA but not as efficiently as PLN. In addition, surprisingly, major differences between a previously published PKAc-WT PLN structure (PDB ID 3O7L) and ours were observed. Our structural models for the three complexes all show a monomeric form of PKAc that forms a 1:1 complex with PLN, which is consistent with the solution behavior of PKAc. In contrast, the 3O7L structure shows PKAc forming a dimer that makes a 2:1 PKAc:PLN complex in the crystal. Further analysis of the 3O7L structure indicates that the dimeric assembly of PKAc is likely an artifact due to crystal packing. The validity of our model is further supported by biophysical and biochemical assays with a series of PLN mutants, designed based on the observed interactions between PLN and PKAc in our structure. Thus, our structure represents only the second physiologically relevant structure of a PKAc-substrate complex, besides the PKAc-ryanodine receptor 2 (RyR2) complex (*37*).

## Results

### Structures of PKAc in complex with WT PLN and PLN R9C

R9C is the most well-known PLN mutation associated with DCM. Despite extensive functional studies, the structural basis of this disease-causing mutation remains elusive. Here we present the crystal structures of PKAc in complex with a peptide corresponding to residues 8-22 of WT PLN, at 2.1 Å resolution (Table 1, Figure 1A), and the R9C variant of this PLN peptide, at 3.4 Å resolution (Table 1, Figure 1B). Adenosine 5’-(β, γ-imido) triphosphate (AMP-PNP), a non-hydrolyzable analog of the ATP co-substrate, is also bound to PKAc in both structures. The peptides contain the phosphorylation site. The WT PLN peptide has previously been shown to be a good model substrate that gets phosphorylated as efficiently as the full-length PLN protein (*38, 39*). The electron densities for the majority of the peptides (corresponding to PLN residues 8-19) and AMP-PNP are well-defined in the structures (Figure 2A, S1). Two Mg^2+^ atoms are observed in the catalytic site, similar to other reported PKA structures (*40*). In both cases, PKAc crystallized in a closed conformation with PLN docked to the large lobe and AMP-PNP bound with the small lobe.

**Table 1.** Data collection and refinement statistics for the PKAc-PLN crystals.

| Crystal | PKAc-WT PLN | PKAc-A11E PLN | PKAc-R9C<br>PLN |
| --- | --- | --- | --- |
| $\lambda$ for data collection (Å) | 0.9795 | 0.9795 | 0.9795 |
| **Data collection** |  |  |  |
| Space group | P 2 21 21 | P 1 21 1 | C 2 2 21 |
| *Cell deminsion* (Å) |  |  |  |
| a, b, c (Å) | 52.57, 70.49,<br>99.03 | 49.56, 69.37,<br>56.24 | 50.91, 104.88,<br>168.10 |
| $\alpha, \beta, \gamma$ (°) | 90.00, 90.00,<br>90.00 | 90.00, 101.97,<br>90.00 | 90.00, 90.00,<br>90.00 |
| Resolution | 42.14-2.16 | 33.12-2.80 | 44.19-3.43 |
| Rmerge† | 0.194 (0.907) | 0.139 (0.527) | 0.165 (0.738) |
| Average I/$\sigma$(I) | 9.9 (1.8) | 8.5 (1.8) | 10.3 (2.5) |
| Completeness (%) | 91.42 (68.22) | 99.11 (94.08) | 96.35 (95.02) |
| Redundancy | 6.9 (6.1) | 3.3 (3.1) | 5.4 (5.7) |
| **Refinement** |  |  |  |
| Resolution (Å) | 42.14-2.16 | 33.12-2.80 | 44.19-3.43 |
| Highest resolution shells (Å) | 2.24 (2.16) | 2.78 (2.83) | 3.56 (3.43) |
| No. of reflections | 18,585 | 9,242 | 6088 |
| Average B-factor | 30.88 | 37.93 | 95.81 |
| Protein | 30.52 | 37.97 | 95.92 |
| Ligands | 24.40 | 34.94 | 86.85 |
| Water | 36.01 | 37.30 | - |
| R<sub>work</sub> | 0.179 (0.230) | 0.203 (0.294) | 0.257(0.339) |
| R<sub>free</sub> | 0.227 (0.290) | 0.252 (0.415) | 0.317 (0.360) |
| RMSD length (Å) | 0.008 | 0.002 | 0.001 |
| RMSD angle (°) | 1.20 | 0.490 | 0.370 |
| **No. of atoms** |  |  |  |
| Protein | 2780 | 2672 | 2775 |
| Ligands | 33 | 33 | 33 |
| Water | 238 | 23 | 0 |
| **Ramachandran plot (%)** |  |  |  |
| Most favored | 95.85 | 95.18 | 94.07 |
| Additionally allowed | 4.15 | 4.82 | 5.64 |

**Fig. 1.**
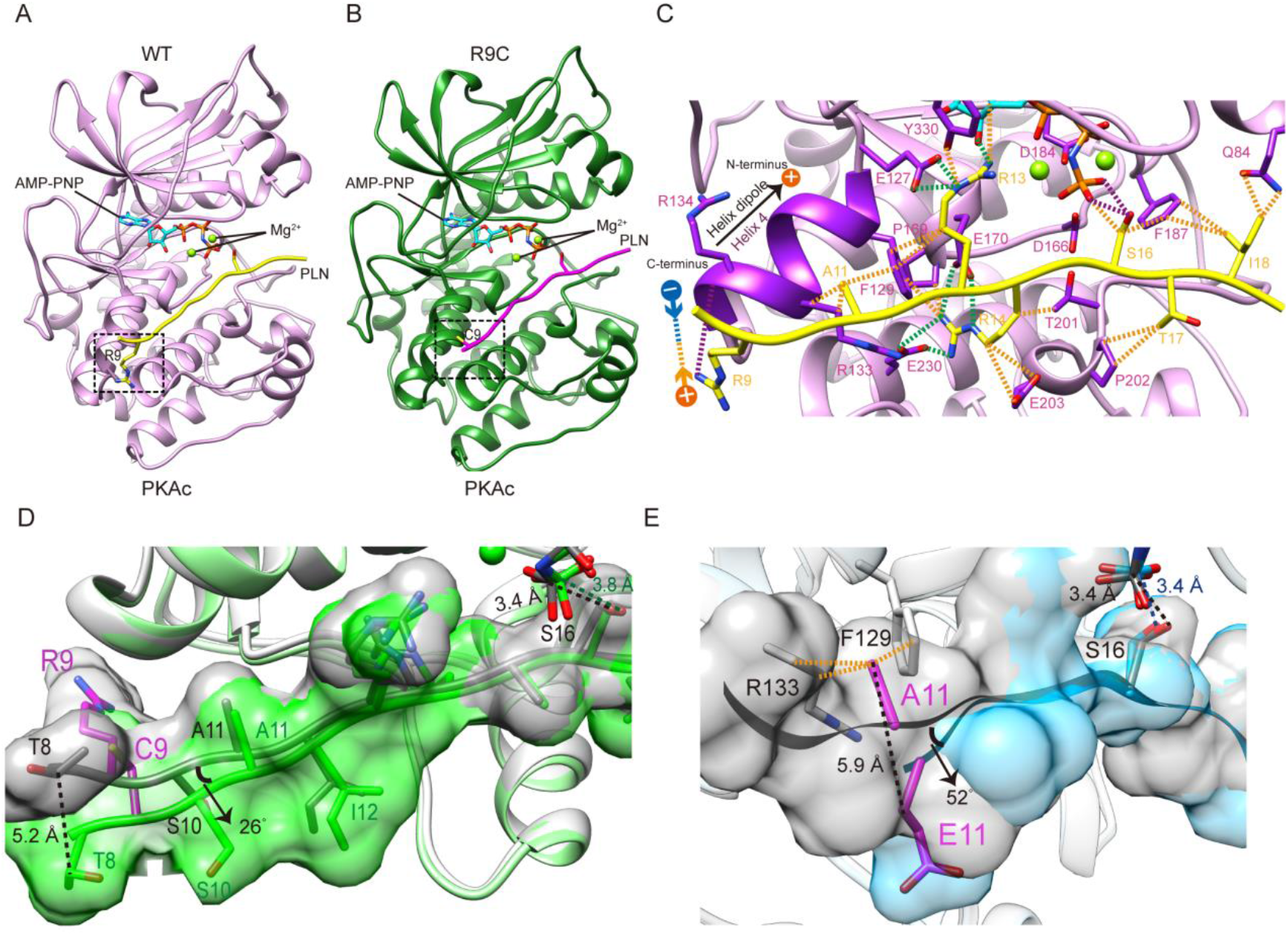
Crystal structures of PKAc-WT/R9C PLN complex. (**A**) Crystal structures of the ternary complex of PKAc, WT PLN, and AMP-PNP. PKA is colored in pink, PLN in yellow, and AMP-PNP in cyan. (**B**) Crystal structures of the ternary complex of PKAc, R9C PLN, and AMP-PNP. PKA is colored in green, PLN in violet red, and AMP-PNP in cyan. (**C**) The interaction between PKAc and WT PLN. The van der Waals contacts (orange), the salt bridges (green), and the hydrogen bonds (purple) are indicated by the dash lines. (**D**) The superposition of PKAc-WT PLN (white-gray) with PKAc-R9C PLN (light green-green). R9C abolishes the electrostatic interaction between Arg9 and the helix dipole of helix 4, inducing conformational changes at the NTR. (**E**) The superposition of PKAc:WT PLN (white-gray) with PKAc:PLN A11E (light blue-cyan). A11E forces the NTR to move away from PKAc without affecting the structure at the catalytic center.

**Fig. 2.**
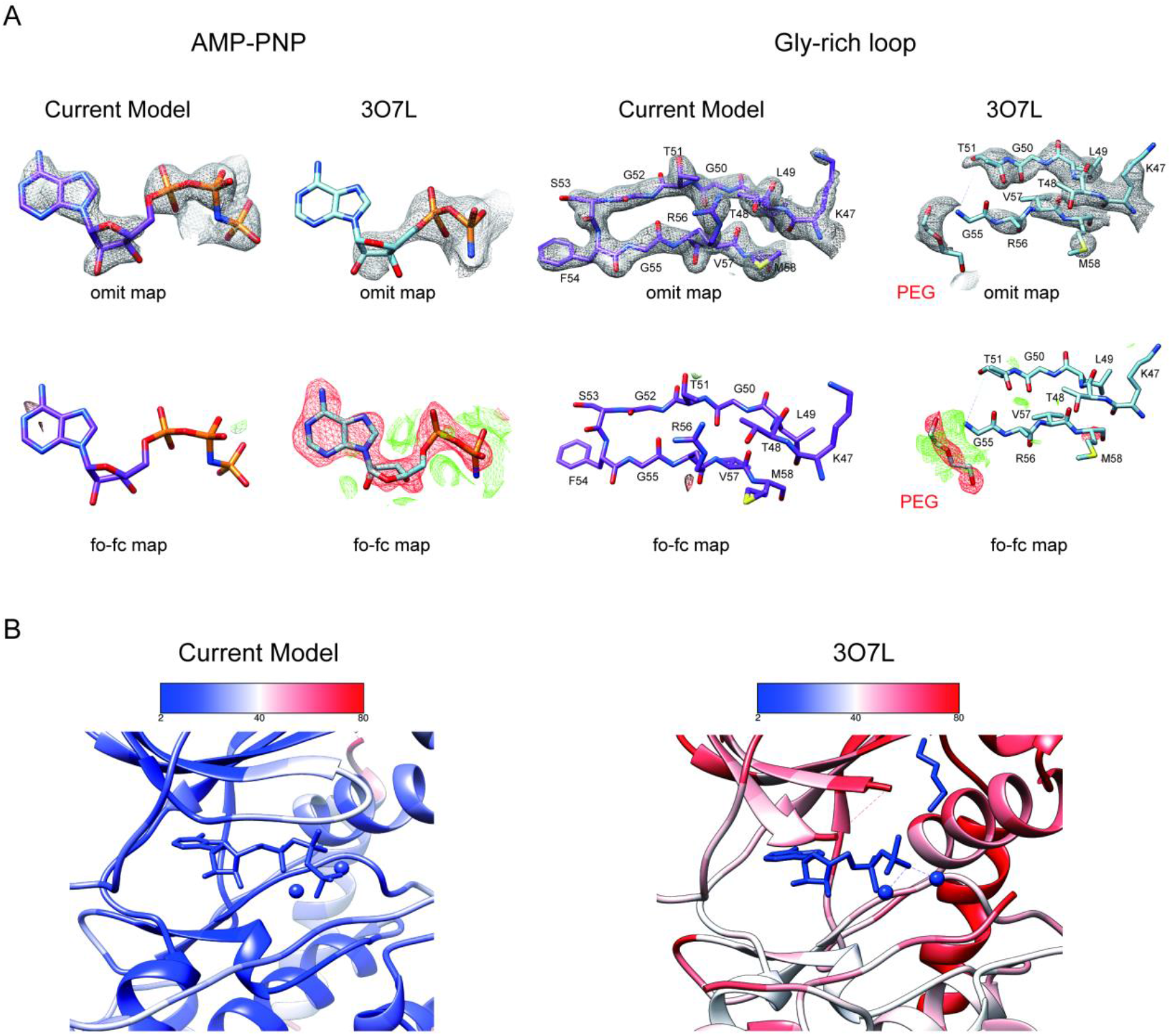
Comparison of two structures of PKAc-PLN complex. (**A**) Comparison of the electron density maps for AMP-PNP region and PKA Gly-rich loop region from our structure (violet red) and 3O7L (blue). The upper panel shows omit maps displayed at contour level of 3σ (AMP-PNP) and 1.5σ (Gly-rich loop), respectively, and the lower panel shows the mFo-DFc difference maps displayed at contour levels of 3σ. (**B**) Zoomed-in view of AMP-PNP binding sites from our structure (left) and 3O7L (right) colored by B-factors.

In the complex of PKAc with WT PLN, the peptide substrate adopts an extended conformation. The N-terminal region (NTR) of WT PLN (Thr8 to Ile12) makes extensive interactions with helix 4 of the large lobe of PKAc. The binding is mainly mediated by an electrostatic interaction between the positively charged side chain of PLN Arg9 and the negative dipole moment of α-helix 4 and by a hydrophobic interaction between Ala11 of PLN and Phe129 of PKAc (Figure 1C). The hydroxyl group of the phosphorylation site, Ser16, is ∼3.4 Å away from the γ-phosphate group of AMP-PNP, similar to other reported PKA structures (*40*). The asymmetric unit (ASU) of our structure contains one PKAc bound to one PLN (Figure 1A).

In the structure of the PKAc complex with PLN R9C, the NTR diverges from the structure of WT PLN bound to the enzyme. The mutation to Cys abolishes the interaction between the positively charged side chain of Arg9 and the negatively charged helix dipole at the C-terminal end of helix 4 from PKAc and a hydrogen-bond between PLN Arg9 and the main chain of PKAc Arg134, which releases the NTR of PLN from binding to the large lobe (Figure 1C, D). R9C turns the NTR counter-clockwise by 26° around the hinge formed by Ala11 and shifts the C_β_ of residue Thr8 by 5.2 Å. The surface area of the enzyme-peptide interface decreases by 62.9 Å^2^, as a consequence of the R9C mutation, which is predicted to strongly diminish the binding of PLN. The structural change in the NTR allosterically affects the conformation of the catalytic center. The C_α_ of Ser16 is shifted by 0.7 Å, which results in a conformational change of the side chain of Ser16, a displacement of AMP-PNP, and a subsequent ∼0.4 Å increase in the distance between the γ-phosphate of AMP-PNP and the hydroxyl group of Ser16 (Figure 1D).

### DCM mutations reduce the binding of PLN and activity of PKA

To test whether the DCM mutations affect the binding between PKAc and the PLN-derived peptide, we characterized their interactions by SPR. To understand the interactions between the three components in the PKAc/AMP-PNP/substrate ternary complex, we first measured the affinity between PKAc and AMP-PNP in the absence of substrate. It shows a dissociation constant (K_D_) around 110 μM (Figure S3A). The binding affinity of WT PLN peptide to PKAc is clearly influenced by AMP-PNP: in the presence of 500 μM AMP-PNP the K_D_ is ∼180 μM, similar to the reported affinities for other PKA substrates such as kemptide and ryanodine receptor (RyR) (*37, 41*), while in the absence of nucleotide, no binding could be detected (Figure 3A, Figure S3B, C). Thus, we chose to include 1 mM of AMP-PNP for all the following SPR experiments involving the formation of ternary complexes.

**Fig. 3.**
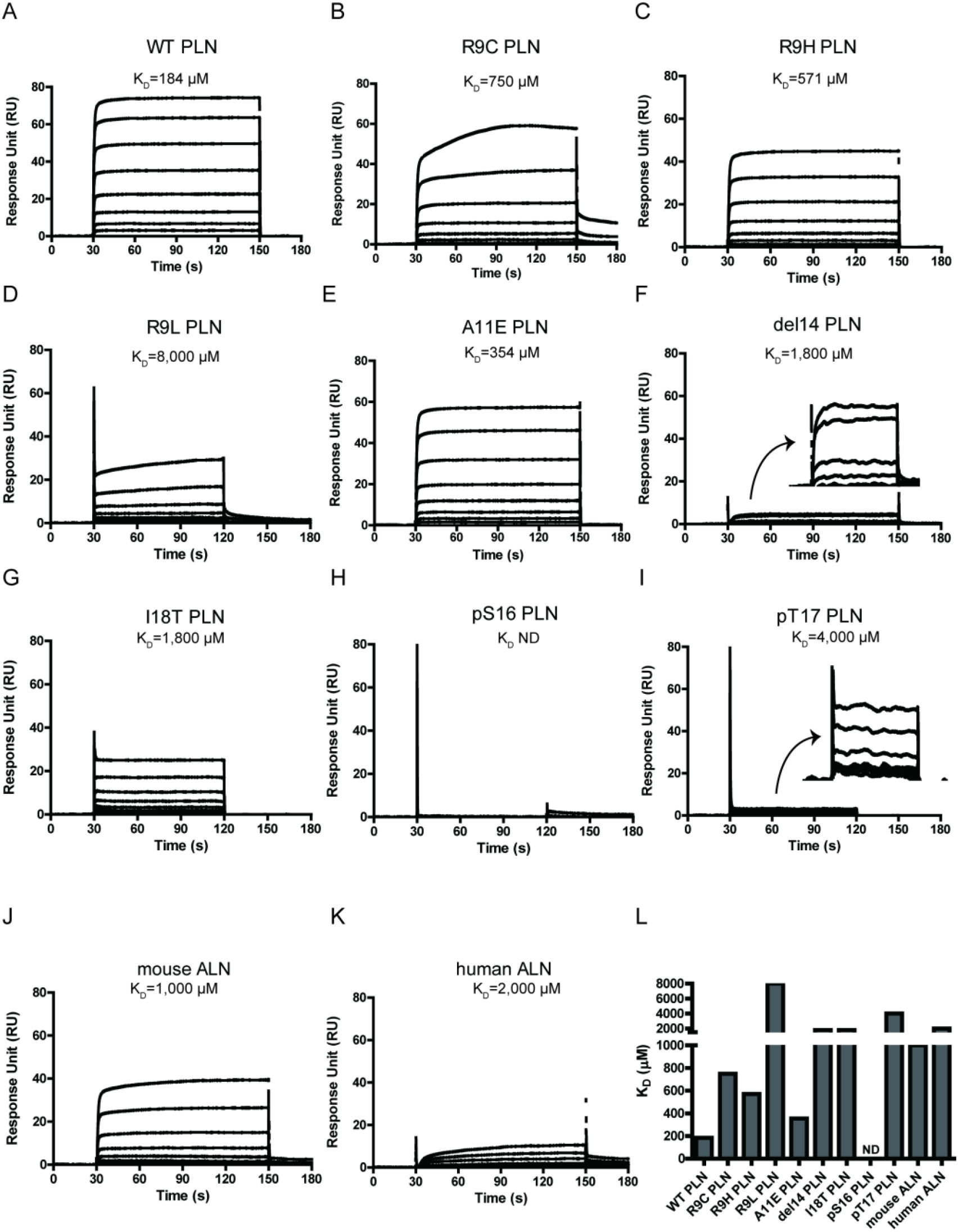
Surface plasmon resonance (SPR) analysis of PKAc-PLN/ALN interactions. (**A-K**) SPR sensorgrams of the binding of the PLN/ALN peptides onto immobilized PKAc. The calculated K_D_ values are displayed above the corresponding sensorgrams. (**L**) The relative K_D_ values of PKAc with different peptide substrates measured by SPR assay.

The K_D_ value of PLN R9C is about four-fold higher than that of WT PLN (Figure 3A, B). This difference confirms that the loss of interactions between the NTR of the peptide and the large lobe of PKAc is linked to a decrease in affinity for PLN R9C. We further tested the impact of four other DCM mutations, including R9H, R9L, ΔR14, and I18T, as well as an artificial mutation A11E, on the interaction between PKAc and PLN. Generally, all of them decrease the binding affinity compared to the WT PLN (Figure 3C-G). Among all the DCM mutations, R9H, the least deleterious of the disease-associated mutations, is the mildest, with an affinity 3.2-fold lower compared to WT. The replacement by histidine partially retains the positive charge at this position and might keep weak contact with the negatively charged helix dipole of helix 4 (Figure 1C). The R9C PLN peptide, which should lack any positive charge character at this position, has a slightly higher K_D_ value compared to the R9H variant. In contrast, the replacement of Arg9 by leucine, which has a non-polar side-chain, shows a much larger weakening effect. Arg14, from the classic R-R-X-S/T motif, forms extensive interactions with PKAc, involving a salt bridge network with Glu170 and Glu230 and van der Waals contacts with Phe129, Thr201, Pro169, and Glu203 of PKAc (Figure 1C), similar to what was seen in previous studies with other peptides known to bind PKAc (*40, 42–48*). Therefore, it is not surprising that the deletion of Arg14 can cause a dramatic 10-fold reduction in binding affinity because it will not only cause the change at Arg14, but also make all residues upstream of residue 14 out of register. The I18T mutation has a similar effect on the affinity as the deletion of Arg14. Ile18 forms extensive van der Waals contacts with Gln84 and Phe187 of PKAc. Replacement by the smaller and more hydrophilic threonine would cause the loss of contacts, which weakens the binding (Figure 1C). PLN A11E exhibits an approximately two-fold elevated K_D_ value, relative to WT PLN. Given the increases in side-chain size and polarity, this mutation likely disrupts the interaction between the methyl group of Ala with Phe129 on α-helix 4 of PKAc, causing the affinity to decrease.

The strength of the interactions between PKAc and substrate peptides was also examined by measuring the thermal stability of the complexes in the solution phase. The addition of WT PLN peptide to PKAc increases its melting temperature (T_m_) by 1.4 °C. In contrast, the mutant PLNs show less contribution to the increase of PKAc thermal stability (Figure 4A, B). Among the five mutations, R9L and ΔR14 show the least stabilizing effects, consistent with the SPR result. Thus, the observed affinity decreases of the DCM-associated peptide variants are not an artifact of PKAc immobilization.

**Fig. 4.**
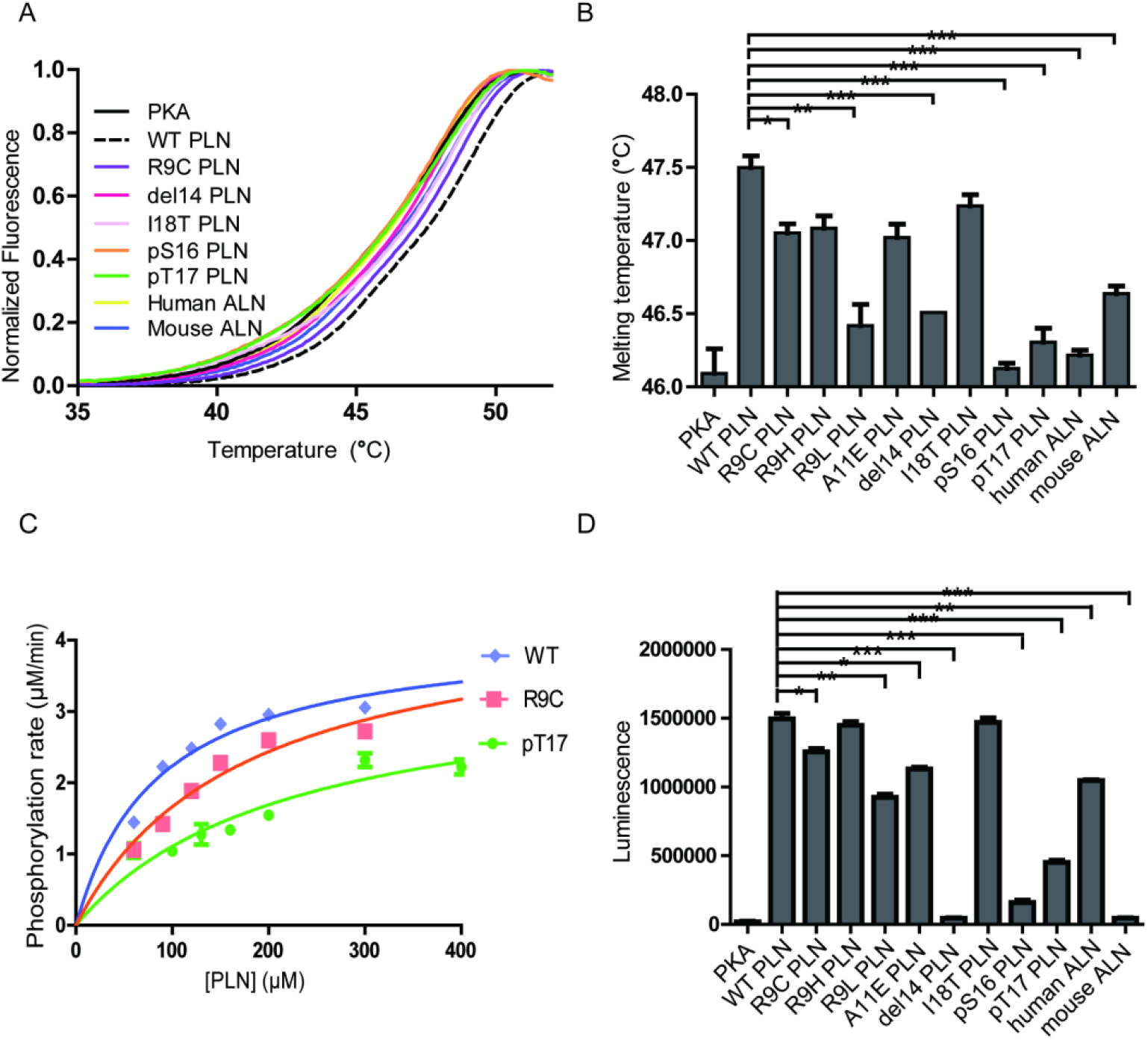
Thermal melt analysis of PKAc-PLN/ALN complexes and the activities of PKAc measured by ADP-Glo assay. (**A**) The averaged thermal melt curves from four replicates each of PKAc complexed with different peptides. (**B**) A bar graph comparing the melting temperatures of PKAc complexed with different peptides. Error bars show the standard deviation. ***P < 0.0001, **P < 0.001, *P < 0.01 (one-tail student T-test). (**C**) A plot of vi vs. [PLN peptide] for PKAc-catalyzed phosphorylation of WT-, R9C-, and pT17-PLN substrates. The data were fit to the Michaelis-Menten equation. (**D**) The relative activities of PKAc with different peptide substrates measured by ADP-Glo assay.

The enzyme kinetic constants of PKAc for WT- and R9C-PLN peptides were determined by an ADP-Glo kinase assay (Figure 4C). The turn-over numbers (*k_cat_*) are generally in the same ballpark with previously reported *k*_cat_ values determined using PLN, kemptide, SP20 and RyR2, as substrates (*37, 38, 49, 50*). The *k_cat_* values for both PLN variants remain relatively unchanged, suggesting that the increased distance and altered orientation between the hydroxyl group of Ser16 in PLN R9C and the gamma-phosphate of AMP-PNP do not significantly impact transition state stabilization during catalysis of phosphate group transfer. The K_M_ value for PLN R9C is 2-fold higher than WT PLN (Table 2, Figure 4C). This difference in K_M_ and K_D_ values is similar for these peptide substrates, which could indicate that the lower catalytic efficiency seen with PLN R9C is mostly due to decreased substrate binding to the enzyme. Our observations suggest that the phosphorylation level of R9C PLN should be lower compared to WT PLN under physiological conditions, which is consistent with previous measurements of their phosphorylation levels in cells made by Western blot (*19, 23, 51*). Next we performed ADP-Glo assays with the other DCM-associated peptide variants, as well as PLN A11E, as substrates at a given concentration near the K_M_ value of WT PLN. Among all mutants tested, R9H has the mildest effect, while ΔR14 shows the largest decrease of PKAc catalytic efficiency (Figure 4D). The enzyme activities of PKAc for different substrates show a roughly similar pattern with their binding affinities determined by our biophysical assays (Figure 4B, 4D), highlighting the importance of the binding affinity of PLN to PKA in DCM disease models.

**Table 2.** Enzyme kinetic parameters for PKAc-catalyzed phosphorylation of different PLN substrates measured by ADP-Glo assay.

|  | WT PLN | PLN R9C | pThr17 PLN |
| --- | --- | --- | --- |
| $V_{\max}$ ($\mu\text{M}/\text{min}$) | $4.1 \pm 0.23$ | $4.5 \pm 0.33$ | $3.6 \pm 0.46$ |
| $K_M$ ($\mu\text{M}$) | $85 \pm 13$ | $173 \pm 25$ | $223 \pm 57$ |
| $k_{cat}$ ($\text{s}^{-1}$) | $6.9 \pm 0.38$ | $7.6 \pm 0.6$ | $6.0 \pm 0.7$ |
| $k_{cat}/K_M$ ($\text{s}^{-1}\text{M}^{-1}$) | $8.1 \times 10^4$ | $4.4 \times 10^4$ | $2.7 \times 10^4$ |

### Comparison of PKAc-PLN structure with a previously reported crystallographic model

Surprisingly, we find that our structure exhibits substantial differences compared to the previously published structure of the complex between PKAc and a peptide corresponding to the first 19 amino acids of human PLN complex structure (PDB ID 3O7L) (Figure 5 A-C, S2G) (*38*). The overall RMSD between 3O7L and our structure of the complex between PKAc and the WT PLN peptide (corresponding to amino acids 8-22 of human PLN) is only ∼0.6 Å, but the RMSD between all modeled Cα atoms of the PLN portions is over 4.4 Å (Figure S4). Another significant difference is that the asymmetric unit (ASU) of our structure contains only one PKAc bound to one WT PLN peptide (Figure 5A). In contrast, the ASU of 3O7L contains two PKAc molecules and one bound PLN whose NTR shows a substantially different conformation and interacts with both PKAc molecules. The second PKAc (Mol B) from the ASU is in a closed non-catalytic conformation. Nontheless, PKAc Mol B makes extensive contacts with the PLN ligand, particularly with the side chains of Tyr6, Leu7, Thr8, Ser10 (Figure 5B). An interface area calculation of 3O7L shows that 29% of the interactions between PLN and PKAc originate from Mol B. In contrast, the interactions of PLN with PKAc in our structure originate mainly from a single PKAc molecule within the same ASU (Figure 5A).

**Fig. 5.**
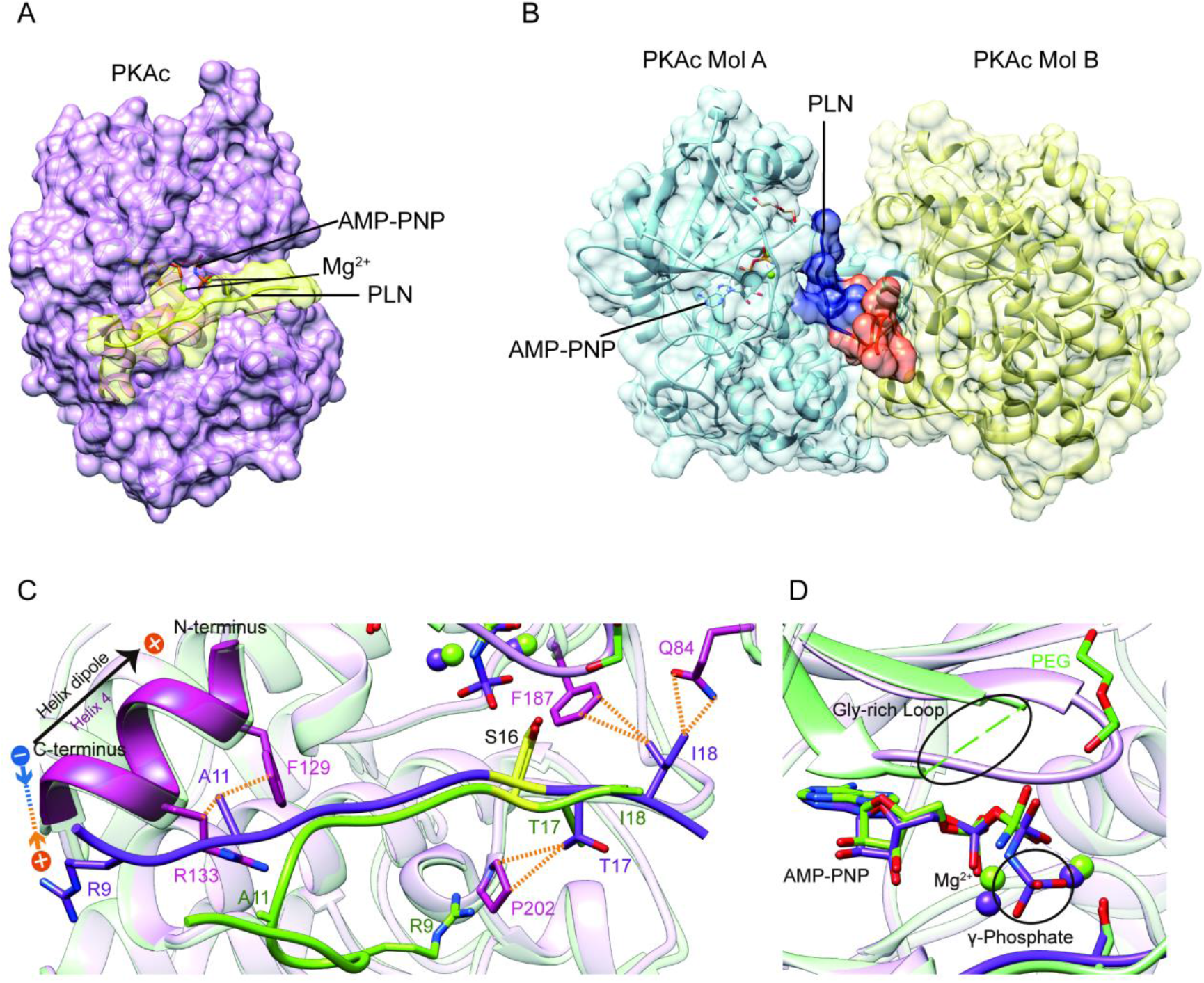
Crystal structures of PKAc-PLN complex. Crystal structures of the ternary complex of PKAc, PLN, and AMP-PNP from the current study (**A**) and a previous study (PDB ID 3O7L)^58^ (**B**). PLN interacts with a single PKAc in our structure and with two PKAc molecules in 3O7L. The NTR of PLN interacts extensively with the second PKAc (Mol B) in 3O7L. (**C, D**) Superposition of our structure (purple) and 3O7L (green) shows that PLNs adopt different conformation at both ends. The γ-phosphate group in AMP-PNP and Gly-rich Loop is missing from 3O7L. The electrostatic interaction between Arg9 and the helix dipole are indicated. The PKA phosphorylation sites are highlighted in yellow.

In order to examine the quaternary structure in solution, purified PKAc complexes were subjected to analytical size-exclusion chromatography. PKAc elutes as expected for a monomer with or without PLN peptide in the presence of AMP-PNP (Figure S2A-C). The monomeric assembly of PKAc:PLN peptide complexes is unaffected by N-terminal truncation, as peptide ligands corresponding to amino acids 1-19 and 8-19 of human PLN eluted similarly. Likewise, the R9C mutation in the PLN sequence did not change the assembly state of the enzyme:peptide complex (Figure S2D, E). Therefore, the interactions that PLN makes with Mol B in the 3O7L structure do not exist in solution but only occur due to the crystal packing. In all other available complex structures of PKAc, there is also only one PKAc molecule bound with one substrate or inhibitor, suggesting the 1:1 ratio should be the common physiological form.

A further difference lies in the active site. Our model shows clear electron density for AMP-PNP and the nearby glycine-rich loop (gly-loop) (Figure 2A, S1). In 3O7L, the γ-phosphate of AMP-PNP was not modeled, and there is a clear negative difference density for the rest of molecule according to the map generated using the previously deposited structure factor data, reflecting a very low occupancy or even an absence of nucleotide (Figure 5D, 2A). In addition, the B-factors for all ligand atoms in 3O7L have been set at a constant value of 2.0, whereas the B-factors for all other atoms from the surrounding regions are clearly much higher, suggesting that AMP-PNP was modeled without any proper refinement (Figure 2B). Further, the 3O7L structural model contains a PEG molecule that is located in a patch of negative difference density, which raises the question of whether or not it is actually present. Neighboring positive difference density is more likely to correspond to the gly-loop according to the comparison of the two structures (Figure 5C,D, 2A, 2B). The gly-loop is right next to the nucleotide and is crucial for nucleotide coordination. Thus, it would be very unusual for a PEG molecule to occupy this loop position, especially when the nearby catalytic loop residues Asp166 and Lys168 are in similar positions in both structures. The observation of a “less-structured” gly-loop in 3O7L could explain the absence of AMP-PNP in that structure.

The last difference is in the C-terminal region of PLN. One of the DCM mutation sites, Ile18 of PLN, shows extensive interactions with Gln84 and Phe187 of PKAc in our model; in contrast, it is mainly facing solvent in 3O7L (Figure 1C). The results from our functional assays (see above) show that the disease mutation I18T clearly reduces the phosphorylation level of PLN and its binding affinity with PKAc, which agrees with our structural model.

### Structure of PKAc in complex with DCM PLN mutant A11E

Residue Ala11 forms hydrophobic interactions with the side chain of Phe129 and the β and γ carbons of Arg133 in our PKAc:WT PLN complex structure; in contrast, the same residue is solvent exposed and is not involved in any interaction with PKAc in 3O7L (Figure 1C). Therefore, to distinguish whether A11E mutation forms contact with PKA or not, we solved the structure of PKAc in complex with the A11E PLN peptide at 2.8 Å resolution (Table 1, Figure 1E). This complex confirms that the mutant PLN can still bind PKAc but with fewer interactions, which explains the decrease in affinity. The mutation flips the side chain of residue 11 and pushes the NTR of PLN away from the large lobe of PKAc (Figure 1E). The C_β_ of Glu11 moves 5.9 Å away from PKAc compared to the C_β_ of Ala11, but the structures of their catalytic centers, including the catalytic loop, gly-loop, two Mg^2+^, and AMP-PNP, are similar. Together with the clear functional effect of A11E (Figure 3E, 4), we propose that Ala11 contributes to the binding of PLN to PKAc as seen in our crystal structure of the PKAc:WT PLN complex.

### Phosphorylation at Ser16 and Thr17 reduce the binding of PLN and activity of PKA

Next, we tested whether phosphorylation at Ser16 (PKA site) and Thr17 (CaMKII site) would affect the binding of PLN with PKAc. The Ser16 side chain points directly to the catalytic center of PKAc. Thus, its phosphorylation would cause steric hindrance with Phe187 and charge repulsion with Asp166, Asp184, and also the gamma-phosphate group from ATP (Figure 1C). This is supported by the previous observation that the K_D_ value of a phospho-serine containing peptide product of PKAc is increased by ∼170 fold compared to the non-phosphorylated substrate (*52*). As expected, we could not detect any significant binding between PLN pSer16 and PKAc by SPR (Figure 3H). In comparison, the Thr17 side chain interacts with the side chain of Pro202 (Figure 1C), so we predict that its phosphorylation would also reduce the binding to PKAc, but to a lesser extent. Indeed, PKAc shows a 20-fold weaker binding towards the phosphorylated Thr17 peptide substrate, but still detectable by SPR, with a K_D_ ∼4 mM (Figure 3I). pSer16 shows a T_m_ value similar to the negative control (PKAc in the absence of PLN), while pThr17 shows a slightly higher T_m_ value (Figure 4A, B), which confirms their low affinities for PKAc found by SPR. The relative activity of PKAc on pThr17 is less than 1/3 of WT PLN, which is mostly due to an increased K_M_ value for this substrate. Only a small residual activity was observed for pSer16, probably due to a small percentage of hydrolyzed pSer16 PLN substrate (Figure 4C, D). The kinetic behaviors of these substrate variants thus reflect their decreased affinities for PKAc.

### Structural dynamics determined by Nuclear Magnetic Resonance (NMR)

To find out how phosphorylation and sequence variations affect the conformation of PLN in the absence of PKA, we solved the structures of peptides corresponding to segments of WT PLN, R9C PLN, pSer16 PLN, and pThr17 PLN by NMR. We analyzed the 20 lowest-energy conformations from all four peptide variants. The WT PLN peptide clearly shows a more dynamic conformation whose structures can be classified into five distinct conformations using Chimera Ensemble Cluster (*53*) (Figure S5A). The dynamic nature of WT PLN can be reflected by the relatively high RMSD value calculated by comparing the representative structures from each ensemble (Figure S5B). In comparison, the R9C, pSer16, and pThr17 PLN variants show a relatively low RMSD among the 20 lowest energy conformations, indicating that these peptides are all less flexible than WT PLN. The structural differences between the R9C-, pSer16-, pThr17-, and WT-PLN peptides might be related to the local charge changes induced by the mutation or phosphorylation, which further affect the intramolecular electrostatic interactions with positively charged Arg13 and Arg14 (Figure S5C). The lower flexibility of R9C PLN and pThr17 PLN might further help to explain their decreased ability to bind PKAc. While none of the conformations of the four peptide variants seem to be significantly pre-organized for binding to the PKA active site, we propose that it might take less energy to rearrange/restructure WT PLN to a proper “bound conformation” before it can be phosphorylated by PKA. If so, both indirect (more energetically costly conformational rearrangement of the peptide during enzyme binding) and direct (loss of a stabilizing electrostatic interaction with the enzyme) effects might contribute to the lower binding affinity (higher K_D_ value) and less efficient conversion to product (higher K_M_ value) of R9C PLN.

### General binding determinant in SERCA-regulating peptides

To study whether other SERCA-regulating peptides can also be phosphorylated by PKA, we tested its activity with another recently identified peptide, called ALN, which is ubiquitously expressed in many tissues. 11AIRRAST17 in human PLN aligns with 14RERRGSF20 in mouse ALN (Figure S6), and both segments contain the R-R-X-S/T PKA recognition motif. As expected, mouse ALN also acts as a PKA substrate, however, PKAc shows about 5-fold lower binding affinity and 1.5-fold lower activity towards mouse ALN compared to human PLN (Figure 3J, 4D). Our PKAc:PLN complex structures show that Ala11 forms a hydrophobic interaction with PKAc, and the replacement of arginine in mouse ALN at this position would introduce charge repulsion with the double arginine at position 133 and 134 of PKAc (Figure 1C). The substitution of Thr17 by the bulky hydrophobic Phe20 in mouse ALN might further cause a clash and reduce the interaction (Figure 1C). We also used human ALN, which lacks the serine phosphorylation site, as a negative control. As predicted, no binding and phosphorylation activity could be detected (Figure 3K, 4B, D).

## Discussion

It is controversial how the mutations in PLN cause DCM. While it is clear that the phosphorylation of PLN by PKA can release its inhibition of SERCA, several models have been proposed to show that the DCM mutations in PLN might change this regulation in either a phosphorylation-dependent or phosphorylation-independent manner (*39, 51, 54-57*). Our data provides structural and functional confirmation that DCM mutations can reduce the binding of PLN substrate to PKA and subsequently its phosphorylation level.

Our work supports a model in which the mutations at positions 9, 14 and 18 of PLN share a common disease mechanism. The cytoplasmic domain of PLN binds with PKAc in a 1:1 ratio through extensive interactions from several key residues, including Arg9, Arg14, and Ile18. The mutations at these three positions have two main effects: 1) change the conformation of the substrate before binding to PKA, as shown by NMR structures; and 2) reduce the binding affinity with PKA, as shown by SPR, thermal melt and ADP-Glo assays, via disruption of enzyme-substrate interactions, as revealed by comparison of the crystal structures of PKAc complexed to WT- and R9C-PLN peptides (Figure S7). Previously it has been proposed that in heterozygous individuals, the aberrant interaction of mutant PLN with PKA may sequester PKA and prevent phosphorylation of WT PLN (*39, 58*). However, our results show that it is not likely that the DCM mutant PLNs can sequester PKA since they interact even more weakly compared to WT PLN. For the structures of PKAc in complex with the two mutant PLNs (R9C and A11E), the positions of the substrates near the PKA catalytic center are relatively conserved, which explains why their turn-over numbers are nearly unchanged relative to WT PLN. Generally, the loss of interactions near the mutation site causes a reduction in ground state affinity and an increase in the K_M_ value, which would result in a decreased phosphorylation level of PLN. Lower phosphorylation levels of PLN in cardiac cells would lead to greater inhibition of SERCA, decreasing heart muscle contractility and relaxation rate. While catalytic efficiency of PKAc with PLN R9C only decreases by ∼2-fold, a corresponding change in phosphorylation level of PLN could be consistent with the relatively mild symptoms of DCM, and even a small increase in SERCA inhibition resulting from such a decrease in PLN phosphorylation would likely compound the Ca^2+^ imbalance in the cell over repeated cycles of cardiac muscle contraction and relaxation. While decreased PLN phosphorylation is likely an important contributor to the physiological dysfunction associated with familial DCM, disease-causing mutations in PLN may have additional consequences, such as altered assembly state of PLN, phosphorylation of PLN by CaMKII, or changes in interactions of PLN with the lipid membrane, that might further increase inhibition of SERCA and act in conjunction with lower PKA-mediated phosphorylation to manifest the disease symptoms.

In addition to changing the interaction with PKAc, mutation or phosphorylation changes the conformational flexibility of free PLN, which might be another reason for the observed decrease in binding. Our NMR results show that R9C-, pSer16-, and pThr17-PLN are generally more rigid compared to WT PLN, probably due to the change in surface charge. Thus, it requires more energy input to reorganize them before binding to PKAc. Previous NMR studies using full-length PLN in the presence of detergents also demonstrated that phosphorylation could change the dynamics of PLN (*59, 60*). A previous study on the intracellular calcium-release channel RyR2 shows that a phosphomimetic at a CaMKII site induces a conformational change from loop to helix, and thus forms a more rigid structure (*37*), similar to the changes in substrate flexibility observed here. However, in that case, the CaMKII site is at a position seven residues upstream of the PKA site. Therefore, the formation of the new helix stabilizes the interaction with PKAc instead of weakening it, as happens with PLN, where the CaMKII site is right next to the PKA site. Subsequently, phosphorylation at the CaMKII site in RyR2 increases the affinity and activity of PKA (*37*), while phosphorylation at the CaMKII site (Thr17) of PLN clearly reduces its ability to be phosphorylated by PKA (Figure S7). Cross-talk between PKA and CaMKII has been reported in a few different cases (*37, 61, 62*). Phosphorylation by one kinase could either facilitate or hinder phosphorylation by a second kinase, and in this way, it connects the signaling networks at different nodes. For PLN, the pThr17-PLN was reported to have the strongest inhibition of SERCA, followed by the pSer16/pThr17 double phosphorylated PLN, while pSer16 had the weakest inhibitory activity (*22*). Thus, the activation of CaMKII on top of the PKA activation could decrease SERCA activity through two related pathways, the reduction of the phosphorylation level on PKA phosphorylation site Ser16 and the weakening of the inhibitory effect of PLN considering pSer16/pThr17 and pThr17 inhibit SERCA more effectively compared to pSer16. It remains to be tested whether phosphorylation by PKA at Ser16 or DCM mutations in PLN also weaken phosphorylation by CaMKII.

So far, three DCM mutations (R9C, R9L and R9H), with different population frequencies, have been identified at the same position on PLN, making Arg9 a DCM mutation hotspot. According to our WT and R9C structures, the replacement of Arg with any neutral amino acid would abolish an important electrostatic interaction between the positively charged arginine and the negatively charged helix dipole and subsequently reduce the phosphorylation level of PLN. The effects of the mutations at the position 9 seems to be correlated to the polarity of the side chain. Histidine, which could be weakly positive, shows the mildest effect, while leucine, which is highly hydrophobic, almost completely abolishes the binding. The effect of cysteine is between the above two replacements. Indeed, it has been shown that R9C and R9L can abolish the inhibition of SERCA, while R9H is more similar to WT PLN (*39, 58*). It requires further investigation to determine whether the clinical severity of these mutations correlates with the change of phosphorylation level.

The previously published crystal structure partially misguided attempts to understand how PKA regulates PLN. There are three clear discrepancies between the previous structure and our structure of the PKAc:WT PLN complex. First, the previous structure shows a sandwich conformation of the complex containing two PKAc and one WT PLN, with the NTR of PLN interacting extensively with the second PKAc molecule in the ASU. This binding mode is clearly a crystallization artifact since the PKAc:WT PLN complex has a monomeric form in solution. Second, the electron density map of 3O7L does not support the presence of AMP-PNP and PEG in their model. Indeed, the difference density is not compatible with a PEG molecule and the area is most likely occupied by the catalytically important gly-loop. Third, the sidechain of DCM mutation site Ile18 was modeled in a truncated form (with only the β-carbon remaining) and in a solvent-facing orientation, which cannot explain the decrease in PKAc binding caused by the DCM mutation I18T. This modeling of the Ile18 side chain could be due to the weak electron density in the previous structure. Our structure shows a different conformation of the Ile18 side chain, which clearly interacts with PKAc. Thus, our explanation for the differences between the two structures is as follows: 3O7L presents a 2:1 (PKAc:PLN) complex structure, where the PLN peptide was trapped between two PKAc molecules from the same ASU, and the crystal contacts force the substrate into an unnatural pose that reduces the binding affinity of the nucleotide; our monomeric structure, which was generated using different crystallization conditions, presents a 1:1 (PKAc:PLN) complex structure, consistent with the native solution behavior and with a fully occupied nucleotide-binding site. These errors in the previous model would certainly impair our understanding of the mechanism by which PKA regulates PLN. Our new structure of the PKAc:WT PLN complex shows clear electron densities in these key regions of the substrate-binding interface and the catalytic center, thus avoiding ambiguity in modeling and providing a more accurate structural template.

To identify general rules for PKA substrate binding, we compared the two available structures of PKAc in complex with their physiological substrates: RyR2 and PLN. As expected, the two PKAc molecules are similar to each other with an overall RMSD of 0.66 Å (Figure 6A). For the substrate, the classic PKA recognition motif “RRXS” shows the highest structural similarity with an RMSD of 0.86 Å, while the NTRs of the substrates show greater divergence (Figure 6A). The interactions observed between Arg13 of PLN and Phe129, Glu170, Glu127, Tyr330 of PKA, and Arg14 of PLN and Glu170, Thr201, Glu203, Pro169, Glu230 of PKA, are conserved between several known PKA substrates (*40, 42–46*). This similarity confirms the importance of the RRXS motif in specific substrate recognition of PKA.

**Fig. 6.**
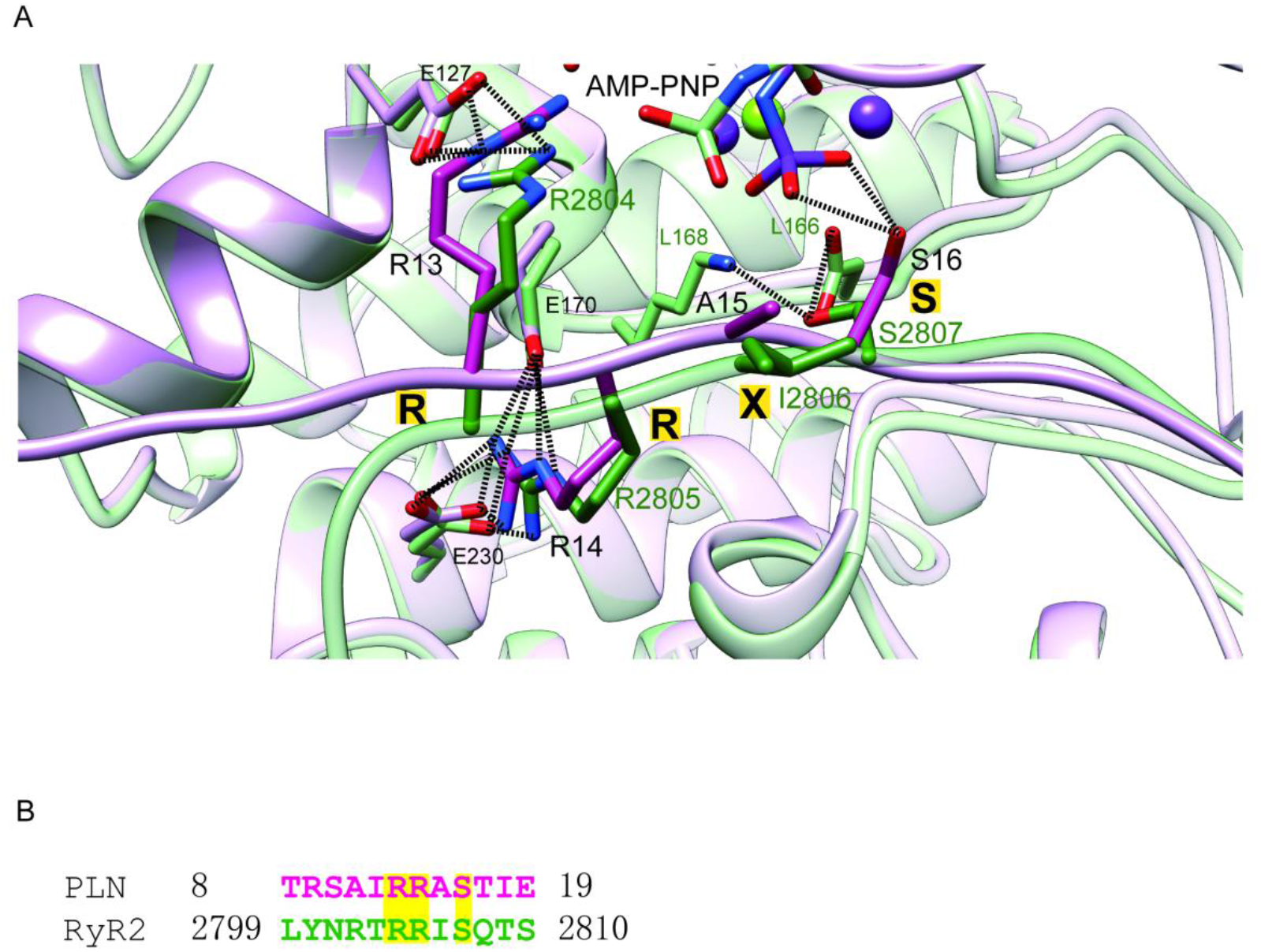
Comparison of PKAc-PLN and PKAc-RyR2. (**A**) Superposition of the crystal structures of PKAc-PLN (violet) and PKAc-RyR2 (green). (**B**) Sequence alignment of PKA-interacting fragments from PLN and RyR2. The conserved RRXS motif is highlighted.

ALN is a newly identified SERCA-regulating peptide that is expressed more ubiquitously than PLN, particularly in the ovary and testis (*34*). ALN has a longer cytoplasmic loop compared to PLN with a predicted PKA recognition motif (Figure S6). Its phosphorylation at Ser19 has been confirmed in liver, pancreas and heart tissue by mass spectrometry (*35, 36*), but the identity of the kinase remained unknown. Using *in vitro* PKA phosphorylation assays, we confirmed that ALN could indeed be phosphorylated by PKAc, although less efficiently compared to PLN. The physiological importance of this regulation remains to be investigated. The phosphorylation of ALN by PKA in mice but not humans could be relevant for understanding animal models of heart disease and how these animal models might behave differently from humans.

## Methods

### Cloning, expression, and purification of protein

The gene encoding PKAc was cloned into the pET-28a-HMT vector, which contains a hexahistidine tag, an MBP fusion protein and a TEV cleavage site at the N-terminus (*63*). For protein production, the plasmid was introduced into *Escherichia coli* BL21 (DE3) cells. Cells were grown at 37 °C with shaking at 250 rpm in 2YT medium supplemented with 50 μg/mL kanamycin. When the OD_600_ reached ∼0.6, protein production was induced with 0.4 mM isopropyl-β-D thiogalactopyranoside (IPTG) and incubated at 18 °C for another 24 h. The cells were harvested by centrifugation at 8000 g for 10 min and disrupted via sonication in lysis buffer (10 mM HEPES pH 7.4, 250 mM KCl, 10 mM BME, 25 mg/mL DNase I, 25 mg/mL lysozyme, 1 mM PMSF). The cell debris was removed by centrifugation at 40,000 g for 30 min. The soluble fraction was filtered through a 0.22-μm filter and loaded onto a 5 mL His Trap HP column (GE Healthcare) pre-equilibrated with buffer A (10 mM HEPES pH 7.4, 250 mM KCl, 10 mM BME). The column was eluted using a linear gradient of 20– 250 mM imidazole in buffer A. The eluted PKAc was cleaved with recombinant TEV protease at 4 °C overnight, followed by purification using an amylose resin column (New England Biolabs) to remove the His-MBP-tag. The samples were loaded to an amylose column pre-equilibrated with buffer A, and eluted with the same buffer plus 10 mM maltose. The flow-through from the amylose column was loaded onto another HisTrap HP column (GE Healthcare) to further remove the fusion tag. PKAc was further purified using a SP Sepharose high-performance column (GE Healthcare) with a linear gradient from 20 to 500 mM KCl in elution buffer (10 mM Tris pH 6.8, 10 mM BME). Finally, the PKAc was concentrated using Amicon concentrators (10 K MWCO from Millipore) and run over a Superdex 200 26/600 gel-filtration column (GE Healthcare) in buffer A. The protein purity was examined by SDS-PAGE with a 15% (w/v) acrylamide gel (Figure S2). The protein sample was concentrated to 10 mg/mL and exchanged to a buffer containing 10 mM HEPES pH 7.4, 50 mM KCl, 10 mM BME for storage at −80 °C.

### Crystallization, data collection, and structure determination

Peptide synthesis of WT and mutant PLN_8-22_ was performed by Genscript Biotech Corporation. The purities of the peptides were > 98% as assessed by analytical HPLC and their molecular masses were verified by ESI-MS. The PKAc:AMP-PNP:PLN_8-22_:Mg^2+^ complex was formed by combining a 1:10:10:10 molar ratio mixture of PKAc (6.5 mg/mL), AMP-PNP, PLN_8-22_ and MgCl_2_ in 10 mM HEPES (pH 7.4), 150 mM KCl, and 10 mM BME at room temperature for 5 min.

Initial crystallization screening was performed by the sitting-drop vapor-diffusion method using commercial crystal sparse matrix screen kits from Hampton Research and Molecular Dimensions. The crystal setting was carried out in 96-well format using a 1:1 ratio with an automated liquid handling robotic system (Gryphon, Art Robbins). After obtaining the initial hits, optimization of crystallization conditions was carried out using hanging-drop vapor-diffusion in a 24-well format. The best crystallization condition for the complex with WT PLN contains 0.1 M BIS-TRIS pH 6.5, and 25% w/v PEG 3350; the best condition for the complex with R9C PLN contains 0.1 M HEPES, pH 7.5, 0.2 M MgCl_2_, and 25% PEG 3350; the best condition for the complex with A11E contains 0.1 M HEPES, pH 7.5, 0.2 M NaCl, and 25% PEG 3350. Crystals were mounted in Cryo-loops (Hampton Research) and flash-cooled in liquid nitrogen with a reservoir solution containing 25% glycerol as cryoprotectant. Diffraction data were collected on BL17U1 at Shanghai Synchrotron Radiation Facility (SSRF) to resolutions of 2.4 Å (PLN_WT_), 3.2 Å (PLN_R9C_), and 2.8 Å (PLN_A11E_), respectively. The dataset was indexed, integrated, and scaled using the HKL3000 suite (*64*). Molecular replacement was performed using the crystal structure of PKAc complexed with a 20-amino acid substrate analog inhibitor as a search model (PDB ID 2CPK) by PHENIX (*65*). After running Phaser-MR, we replaced the model sequence with the object sequences. The structure was further manually built into the modified experimental electron density using Coot (*66*) and refined in PHENIX^57^ in iterative cycles. The data collection and final refinement statistics are shown in Table 1. All structure figures were generated using UCSF Chimera (*53*).

### Fluorescence-based thermal shift assays

The protein melting curves were measured using a fluorescence-based thermal shift assay (*67*). The Sypro orange dye (2×), PKAc (0.2 mg/mL), AMP-PNP (500 μM), and a PLN peptide variant (1 mM) were mixed in 8 strip tubes (Axygen). The tubes were then transferred to a centrifuge and rotated to remove any bubbles and homogenize the system. The tubes were then placed into a Quant Studio 6 Flex real-time PCR machine (Life). The temperature was increased from 10 °C to 95 °C with a ramping rate of 0.033 °C/s. All measurements were performed in triplicate. The melting temperatures were obtained by taking the midpoint of each transition.

### ADP-Glo kinase assay

The kinase activity of PKAc was measured using the ADP-Glo kinase kit (V9101; Promega) according to manufacturer’s instructions. Phosphorylation of PLN peptides were performed at 30 °C for 30 min in 50 μL kinase buffer (10 mM HEPES pH 7.4, 150 mM KCl, 20 mM MgCl_2_, 2 mM DTT) supplemented with 200 μM ATP, 10 nM PKAc and 90 μM peptide substrates. 25 μL samples were removed and terminated by adding 25 μL ADP-Glo™ reagent followed by incubation at room temperature for 40 min. Kinase detection reagent was prepared by combining kinase detection buffer with kinase detection substrate based on the manufacturer’s instructions. 50 μL kinase detection reagent was added and incubated at room temperature for 40 min to convert ADP to ATP. The luminescence signal was read by a Tecan Infinite M200 Pro plate reader. All measurements were performed in triplicate.

### Surface Plasmon Resonance (SPR) analysis

SPR experiments were carried out to characterize the interaction between PKAc and substrate peptides using a Biacore T200 instrument (GE Healthcare). PKAc was immobilized via standard N-hydroxysuccinimide (NHS) / 1-ethyl-3-(3-dimethylaminopropyl) carbodiimide hydrochloride (EDC) amine coupling on a CM5 (carboxyl methyl dextran) sensor chip (GE Healthcare). Before covalent immobilization of PKAc, the sensor surface was activated by a mixed solution of 0.4 M EDC and 0.1 M NHS (1:1) for 7 min at a flow rate of 10 μL/min. The purified PKAc protein was diluted to 35 µg/mL in 200 µL of immobilization buffer (10 mM sodium acetate, pH 5.5) and immobilized on the sensor chip to a level of 7000 response units (RU). Interactions between PKAc and substrate peptides were monitored by injecting various concentrations of peptides (two-fold serial dilutions starting from 1 mM or 2 mM) in the running buffer containing 10 mM HEPES, pH 7.4, 150 mM KCl, 20 mM MgCl_2_, 1 mM AMP-PNP and 0.005% (v/v) Surfactant P20 at a flow rate of 30 μL/min for 120 s. Dissociation was performed by running the buffer without peptides at the rate of 30 μL/min for 120 s. The RU was obtained by subtracting a control for unspecific binding (the signal from a blank flow cell without PKAc subunit).

### NMR

The PLN peptides were dissolved in 10% or 100% D_2_O. ROESY and TOCSY spectra were recorded at 298K using an 850 MHz Bruker Avance NMR spectrometer equipped with a 5-mm cryogenic probe. NMR spectra were processed using NMRPipe (*68*) and analyzed using NMRFAM-Sparky (*69*). Distance constraints obtained from the assigned NOEs were divided into three classes based on the intensities of NOE cross-peaks: (1) strong: 1.8 Å < d < 2.8 Å; (2) medium: 1.8 Å< d < 3.4 Å; and (3) weak: 1.8 Å < d < 5.5 Å. The solution structure was calculated with cyana 2.1 (*70*). 20 conformers from a total of 100 calculated ensembles with the lowest energy were selected for analysis.

## Conflict of Interests

The authors declare that they have no conflict of interest.

## Acknowledgments

We thank J. Xu from the Instrument Analytical Center of the School of Pharmaceutical Science and Technology at Tianjin University for assisting in using the in-house X-ray diffraction machine, J. Shen at the Tianjin Institute of Industrial Biotechnology Chinese Academy of Sciences for assisting SPR analysis, and the staff at the beamline BL17U1 at Shanghai Synchrotron Radiation Facility.

## Funding

Funding for this research was provided by the National Natural Science Foundation of China (no. 3202207 and 31972287, to Z.Y.), the Natural Science Foundation of Tianjin (no. 19JCYBJC24500, to Z.Y.), CIHR (no. PJT-159601, to F.V.P.), and fellowships from the CIHR and Michael Smith Foundation for Health Research (to O.H.G.).

## Author contributions

Z.Y. and J.Q. conceived of the project. J.Q. solved the crystal structures of PKAc-PLNs and carried out the biophysical analysis. L.L participated in the crystallographic study. J.Z. and Z.L. carried out the NMR analysis. J.Q. led the manuscript preparation with guidance from Z.Y., K.J.W., F.V.P., Y.Z., and O.H.G. Z.Y. supervised the project.

## Data and materials availability

The atomic coordinates and structure factors have been deposited in the Protein Data Bank: PKAc-WT PLN (PDB 7E0Z); PKAc-PLN R9C (PDB 7E11); PKAc-PLN A11E (PDB 7E12).

---

**Fig. S1.**
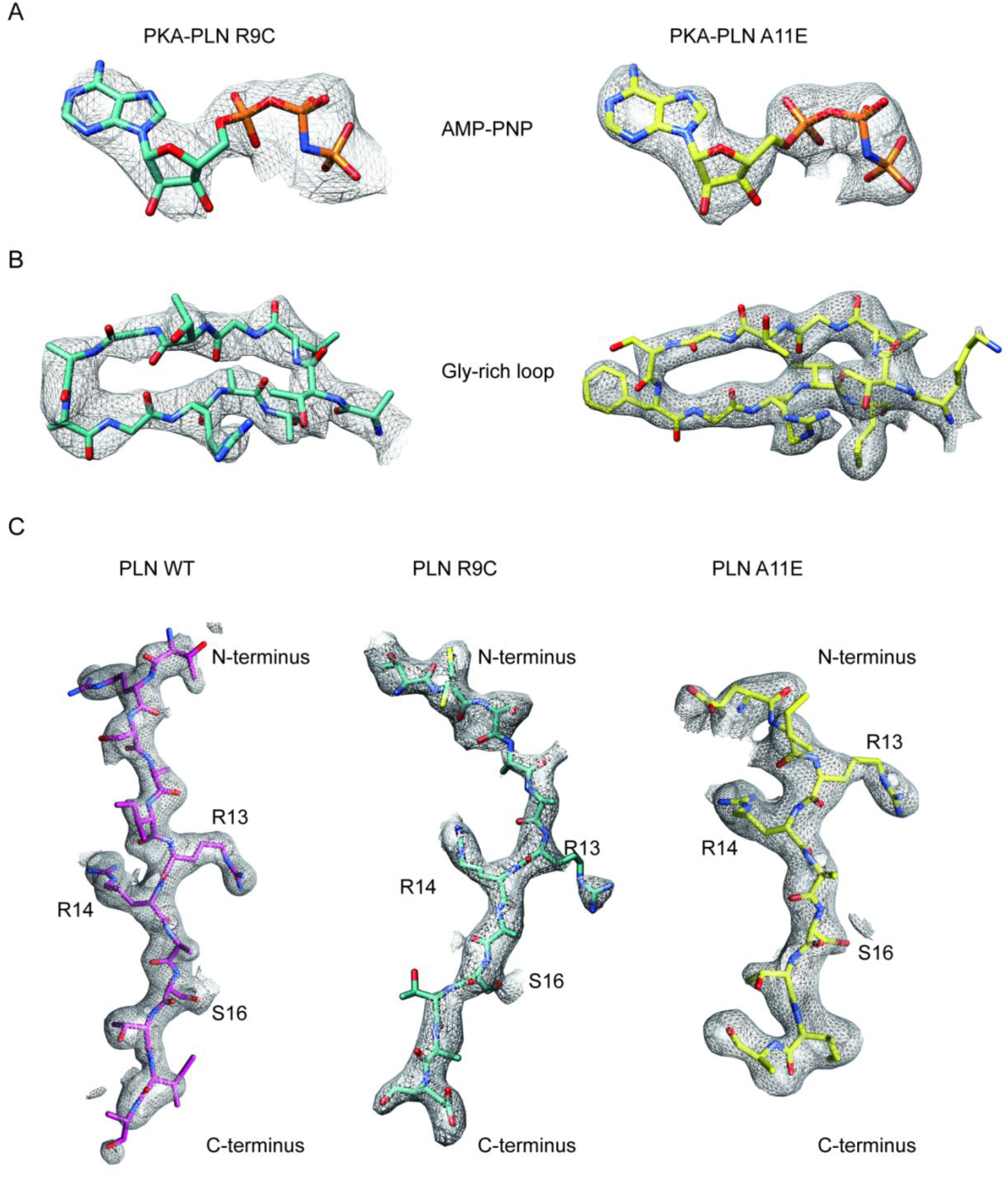
Electron density maps. Density maps for the AMP-PNP region **(A)**, PKAc Gly-rich loop region **(B)**, and PLN peptides **(C)** from the crystal structures of PKAc-PLNs.

**Fig. S2.**
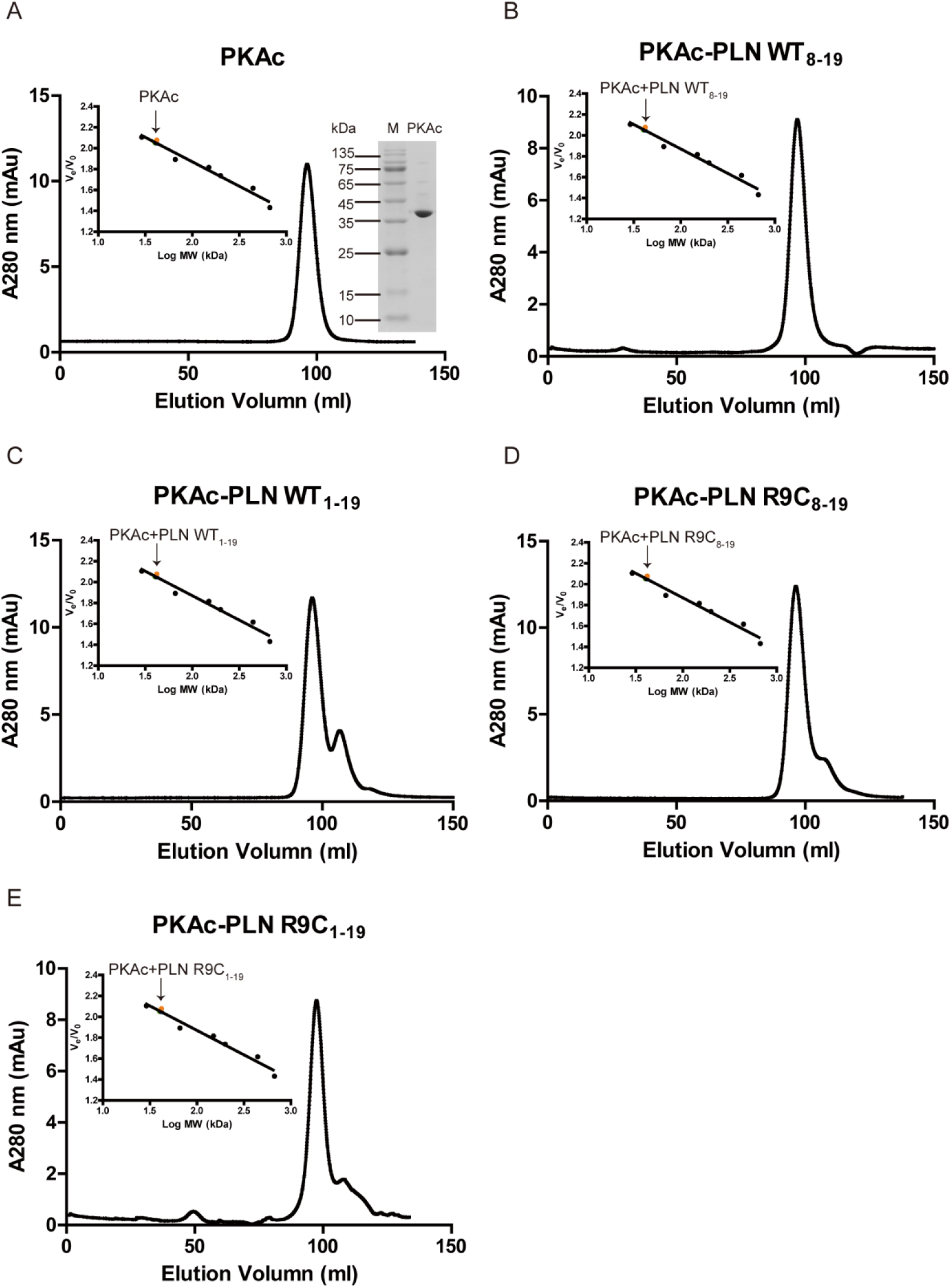
Oligomerization states of PKAc. (A-E) Elution profiles of PKAc (A) or PKAc-PLN complexes (B-E) filtration chromatography using a Superdex 200 16/600 column (GE Healthcare, USA). The right inset in A is a 15% SDS-PAGE gel of purified PKAc. The left inset in A-E shows the plotted standard curve for this column. The molecular weight (MW) estimated from their elution volumes are ∼40 kDa, suggesting monomeric forms in solution.

**Fig. S3.**
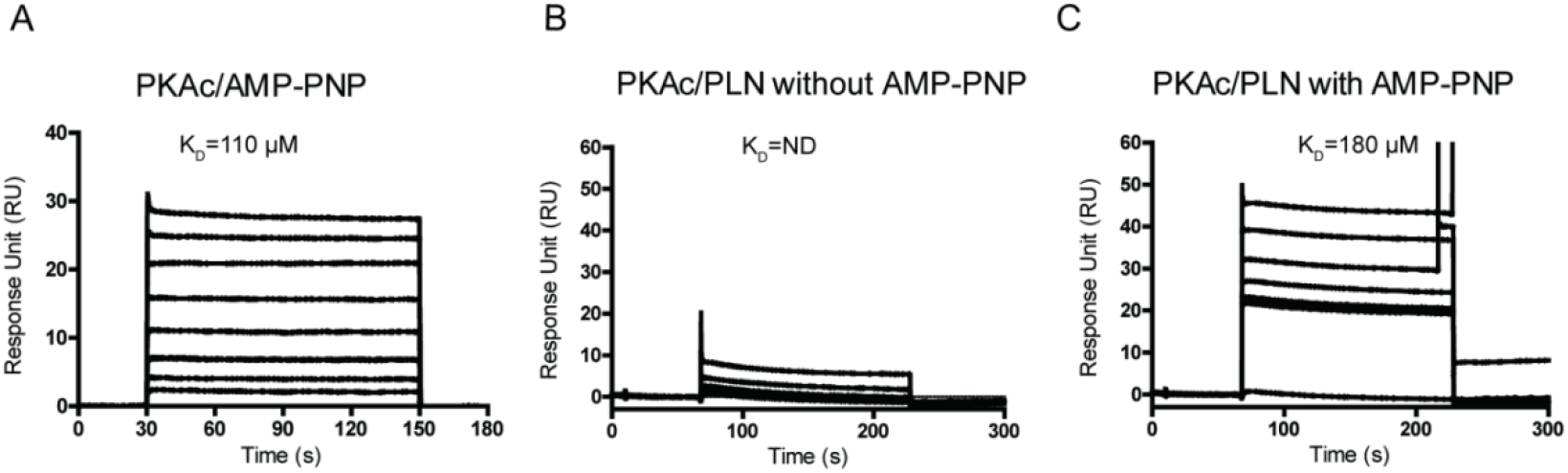
The SPR analysis of PKAc/AMP-PNP interactions. **(A)** SPR sensorgrams for the binding of AMP-PNP with immobilized PKAc. **(B)** and **(C)** SPR sensorgrams for the binding of WT PLN with immobilized PKAc in the absence (B) or presence (C) of AMP-PNP.

**Fig. S4.**
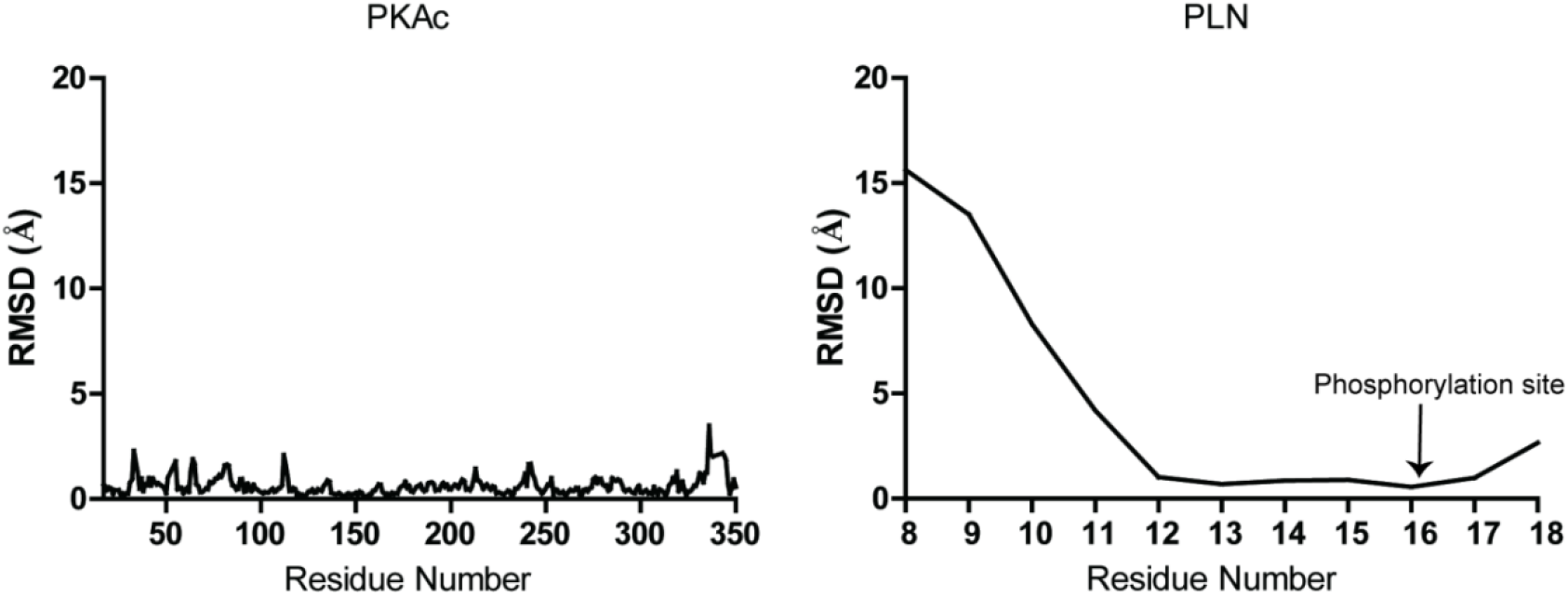
Root mean square deviation (RMSD) Plots. Plots showing per residue RMSD for the PKAc (left) and PLN (right) from our structure relative to 3O7L.

**Fig. S5.**
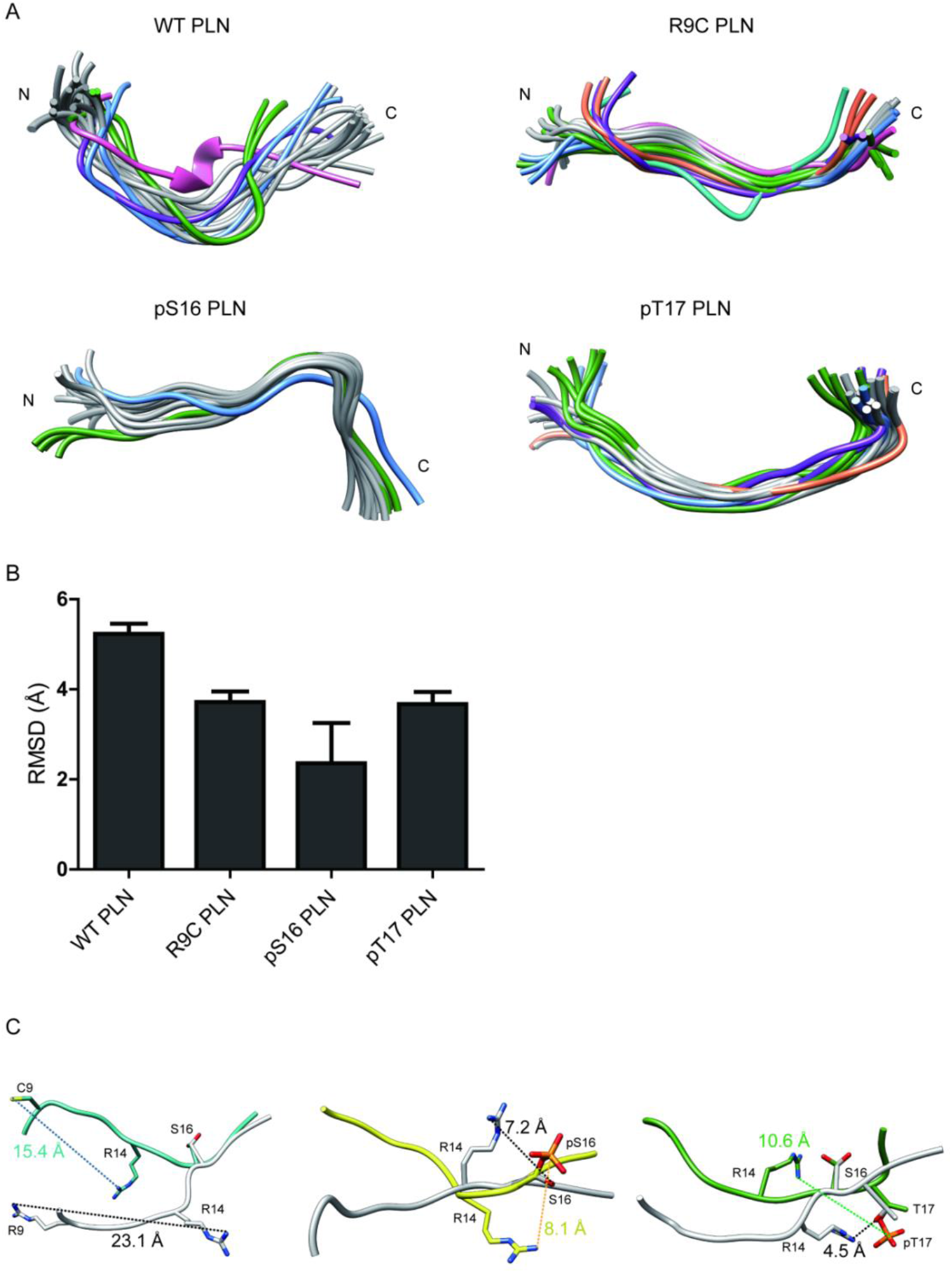
NMR structures of PLN/ALN peptides in the absence of PKAc. **(A)** Ensemble cluster views of the superposition of the top 20 lowest energy structures of WT PLN, R9C PLN, pS16 PLN, and pT17 PLN. The top seven ranked clusters are colored in gray, forest green, cornflower blue, purple, sandy brown, light sea green, and hot pink, respectively. **(B)** The Root Mean Square Deviation (RMSD) between the representative structures from each ensemble cluster. WT PLN clearly shows a higher RMSD than the other four peptides, suggesting its higher flexibility. **(C)** A comparison of the top structures of WT PLN (white) with the ones from R9C PLN (light sea green), pS16 PLN (yellow), pT17 PLN PLN (forest green). S16 is used for the superposition.

**Fig. S6.**
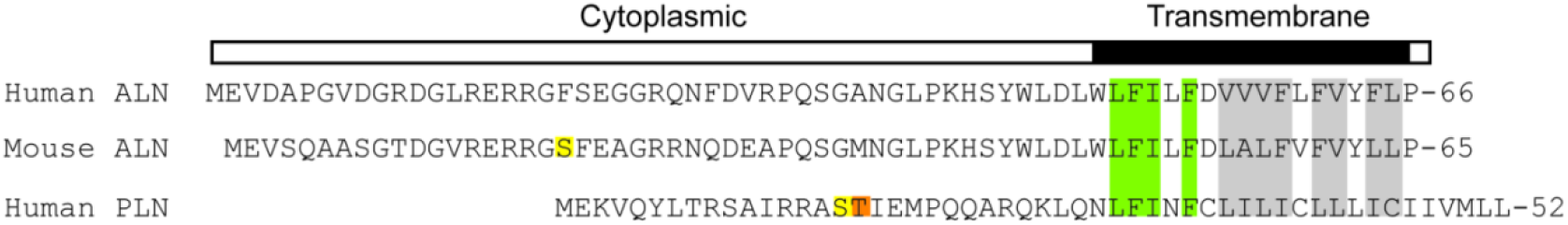
Sequence alignment between PLN and ALN. Identically conserved residues are coloured in green, and weakly similar residues are coloured in grey. The PKA/CaMKII phosphorylation site is colored in yellow and orange, respectively.

**Fig. S7.**
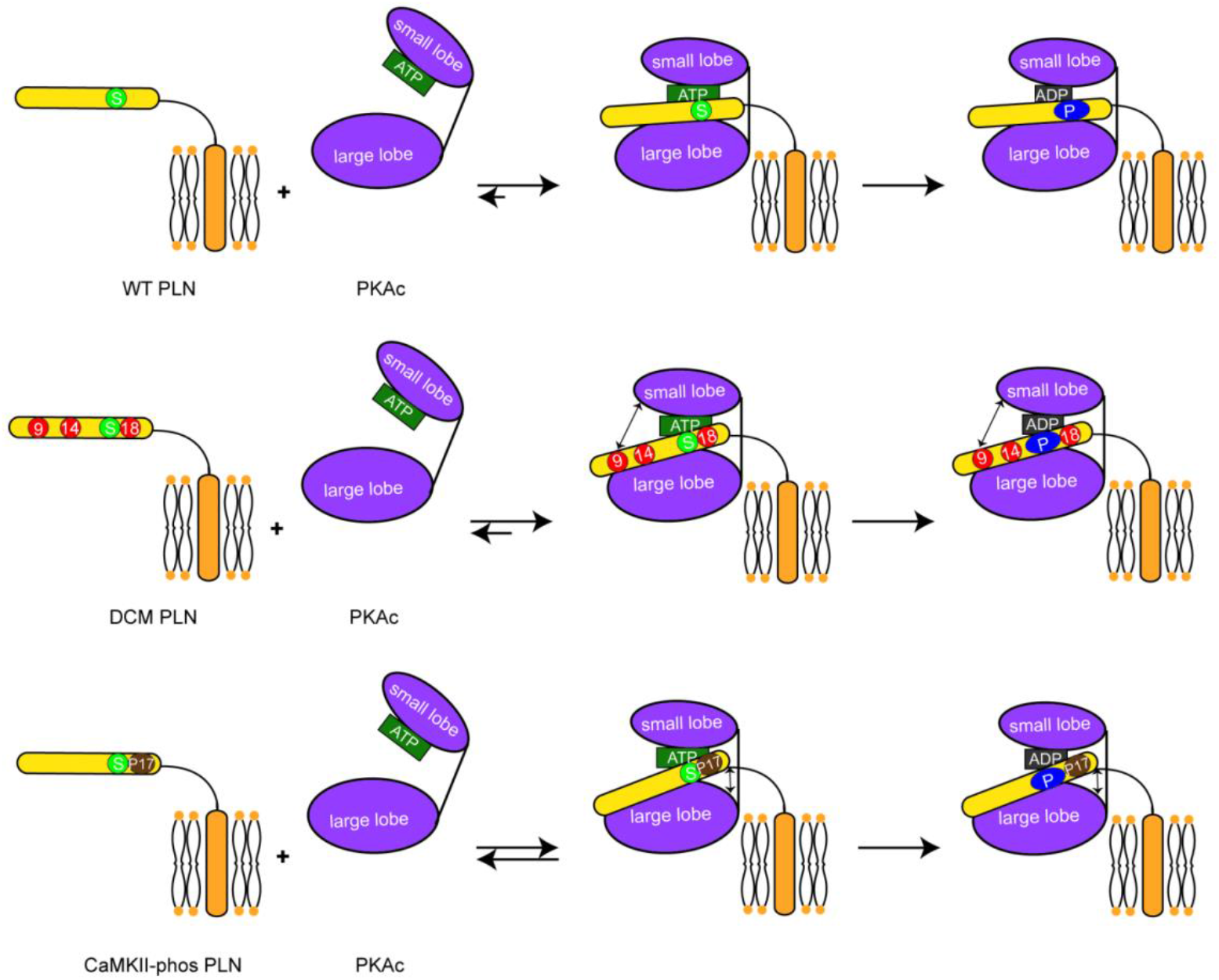
Schematic representation showing the impacts of DCM mutations and CaMKII phosphorylation on the phosphorylation of PLN by PKA. Both DCM mutations at the position 9, 14 and 18 (red circles) and the CaMKII phosphorylation (brown circle) reduce the interaction between PLN (yellow) and PKAc (purple), and subsequently decrease the phosphorylation level of PLN.

## Notes

### Competing Interest Statement

The authors have declared no competing interest.

